# Transcriptome profiling of ulcerative colitis mouse model suggests biomarkers and therapeutic targets for human colitis

**DOI:** 10.1101/2020.08.12.225458

**Authors:** Reza Yarani, Oana Palasca, Nadezhda T. Doncheva, Christian Anthon, Bartosz Pilecki, Cecilie A. S. Svane, Aashiq H. Mirza, Thomas Litman, Uffe Holmskov, Claus Heiner Bang-Berthelsen, Mogens Vilien, Lars J. Jensen, Jan Gorodkin, Flemming Pociot

## Abstract

1.

**BACKGROUND & AIMS:** Ulcerative colitis (UC) is an inflammatory bowel disorder with unknown etiology. Given its complex nature, *in vivo* studies to investigate its pathophysiology is vital. Animal models play an important role in molecular profiling necessary to pinpoint mechanisms that contribute to human disease. Thus, we aim to identify common conserved gene expression signatures and differentially regulated pathways between human UC and a mouse model hereof, which can be used to identify UC patients from healthy individuals and to suggest novel treatment targets and biomarker candidates.

**METHODS:** Therefore, we performed high-throughput total and small RNA sequencing to comprehensively characterize the transcriptome landscape of the most widely used UC mouse model, the dextran sodium sulfate (DSS) model. We used this data in conjunction with publicly available human UC transcriptome data to compare gene expression profiles and pathways.

**RESULTS:** We identified differentially regulated protein-coding genes, long non-coding RNAs and microRNAs from colon and blood of UC mice and further characterized the involved pathways and biological processes through which these genes may contribute to disease development and progression. By integrating human and mouse UC datasets, we suggest a set of 51 differentially regulated genes in UC colon and blood that may improve molecular phenotyping, aid in treatment decisions, drug discovery and the design of clinical trials.

**CONCLUSION:** Global transcriptome analysis of the DSS-UC mouse model supports its use as an efficient high-throughput tool to discover new targets for therapeutic and diagnostic applications in human UC through identifying relationships between gene expression and disease phenotype.

## 2. Introduction

Ulcerative colitis (UC) is an inflammatory bowel disorder that mainly affects the large intestine. Most often, UC initiates from the rectum and affects the mucosal lining and is limited to the innermost layers of the large intestine. Multiple pathogenic factors, including numerous susceptibility gene variants, environmental factors, changes in the gut microbiota and a dysregulated immune response are all associated with UC. Despite this recognition and the identification of apparently relevant factors, a complete understanding of UC pathogenesis is still out of reach and, thus, treatment may not be optimal. An important reason for this unsatisfactory situation is the currently limited comprehension of what the truly relevant components of UC immunopathogenesis are. Thus, given the complex nature of UC, the study of animal models is crucial. A big challenge in using animal models for UC is whether the model’s genes and molecular pathways are analogous to those in humans. Although many coding and non-coding genes between humans and mice are highly similar,^1^ they might be regulated differently, and also the pathophysiology may be different. Expression of both differentially regulated coding and non-coding genes contributes to disease pathology.^2, 3^ Differentially regulated long non-coding RNAs (lncRNAs) and microRNAs (miRNAs) are linked to the change in the expression of many protein coding genes (PCGs) and subsequently, development of disease. Many studies are made without taking the non-coding RNAs, i.e. disease related pathways and differentially expressed genes, into account and thereby, fundamental and essential players in the disease regulatory networks may be overlooked.^4^

Preclinical murine models have been used for *in vivo* assessments of UC, and one of the best established UC models is the dextran sulfate sodium (DSS) model.^5, 6^ DSS is a negatively charged polysaccharide with colitogenic properties that induces colitis when given orally.^5^ Although the mechanisms by which DSS induces intestinal inflammation are not fully understood,^7^ it seems that the exposed colonic monolayer of epithelial cells in the large intestine, specifically the distal colon are being progressively eroded and the lamina propria barrier integrity compromised, in which an enormous number of microorganisms live. Therefore, inflammation is triggered following bacteria (and their products) translocation across the damaged intestinal wall. To our knowledge, a comprehensive investigation of the coding and non-coding colonic and blood transcriptome of the DSS model and its comparison with human UC has not been performed.

We therefore set out to characterize the transcriptomic landscape of the DSS-UC mouse model by performing transcriptomic profiling of PCGs, lncRNAs and miRNAs from colon and whole blood using deep RNA sequencing. We identified differentially expressed lncRNAs and underscored their functional importance by highlighting their functional annotation through the expressed neighboring PCGs. Then we identified the targets for the differentially regulated miRNAs and identified biological processes that these targets are affecting. Further, we looked at the underlying molecular pathways and networks and assessed the similarity between the experimental model and human UC. Our results identified widespread differential regulation of coding and non-coding genes both in the mouse model and human UC with marked similarities in both tissues specifically in colon. Altogether, this study offers a great opportunity to further investigate the role of novel and previously identified differentially regulated genes in UC and to improve molecular phenotyping, which may have marked effects on treatment decisions, drug discovery and the design of clinical trials.

## 3. Materials & Methods

### 3.1. Mouse model, RNA sequencing, data processing and differential expression analysis

Ulcerative colitis was induced using dextran sulfate sodium salt (DSS) in C57BL/6 male mice. The control group was time-matched and received the same drinking water without DSS. Disease activity index (DAI) was recorded. RNA was extracted from colon and whole blood, and total and small RNA were sequenced using the Illumina HiSeq 4000 system. Gene-level quantification corresponding to the total RNA-Seq data was obtained using Stringtie,^8^ and mature miRNA expression was quantified using miRDeep2.^9^ Differential expression was performed using DESeq2.^10^ (Supplementary Methods). The raw total and small RNA-Seq data have been deposited in the Gene Expression Omnibus (GEO) database with accession number GSE155303.

### 3.2. Functional enrichment and network analysis of differentially expressed genes

Functional enrichment analysis on the significantly differentially expressed (SDE) genes (FC > 2, padj≤0.05) was performed using stringApp^11^ in Cytoscape^12^ as well as Ingenuity pathway analysis (IPA) software.^13^

To suggest a functional role for SDE lncRNAs, we performed enrichment on their genomic neighbor genes situated within a span of 100 kb upstream and downstream of the lncRNA. For SDE miRNAs (padj≤0.05), we retrieved a set of miRNA-target genes and performed enrichment on them (Supplementary Methods).

stringApp was used to retrieve STRING networks^14^ in Cytoscape for the set of SDE PCGs in mouse colon and blood and for the SDE PCGs conserved in humans. In order to rank the SDE genes common between mouse and human colon and blood based on their importance, weighted centrality analysis was performed on their STRING network (Supplementary Methods).

### 3.3. Comparison between mouse and human data

In order to compare our mouse data with data from UC patients, we obtained and processed data from 8 different publicly available studies.^15–21^ A set of SDE genes in colon and blood was obtained by combining the different datasets, requiring each gene to be consistently SDE (padj≤0.05) in at least two datasets. For the small RNA-Seq we only used two datasets and required that each miRNA is SDE (padj≤0.05) in at least one of these datasets. The final SDE human lists were intersected with the mouse SDE blood and colon gene sets (padj≤0.05) using orthology relationships extracted from Ensembl (release 97),^22^ and HCOP database^23, 24^ for lncRNAs (Supplementary Methods).

### 3.4. Validation of SDE genes by quantitative real-time PCR

The expression of 19 selected SDE genes common in both mouse and human colon and blood was validated by quantitative real-time PCR (qRT-PCR). For validation, colon and blood cDNAs from 5 DSS-UC mice, 5 control mice, 10 UC patients and 6 controls were used. Detailed information on patient demographics and primer sequences used for qRT-PCR can be found in Supplementary File 1 and procedure detail in the supplementary section. Expression of each gene tested was represented as a FC using the 2^−ΔΔCT^ method. GUSB, B2M, ACTB and TBP were used as the reference genes for normalization.

## 4. Results

### 4.1. Transcriptional profile of UC mouse model

Successful colitis induction in mice was confirmed by combined DAI score, measuring inflammatory markers, histological and micro positron emission tomography imaging analysis (Supplementary Figure 1). For the total RNA-Seq, 40-66M paired-end reads were obtained per sample, with 39-58M in colon and 17-33M in blood, uniquely mapping to the mouse genome. For the small RNA-Seq after cleaning, 15-25M reads were obtained, where 14-20M reads mapped to the set of mouse miRNAs annotated in miRBAse v.22. Principal component analysis (Supplementary Figure 2) shows a clear separation of the disease and control groups on the first principal component in total RNA-Seq colon and blood, as well as small RNA-Seq colon. The separation between the groups is strongly supported by the differential gene expression pattern, suggesting that global gene expression differences can be clearly detected between UC and healthy controls (Table 1 and Supplementary File 2).

**Table 1:** Numbers of SDE genes identified in colon and blood of DSS-UC mice versus control. (For PCGs and lncRNAs |log2FC| > 1, padj≤0.05 and for miRNAs FC > 1.5, padj≤0.05).

|  | Colon | Blood |
| --- | --- | --- |
| PCGs | 2479 ( ▲ 1897 , ▼ 582 ) | 518 ( ▲ 500 , ▼ 18 ) |
| lncRNAs | 282 ( ▲ 135 , ▼ 147 ) | 44 ( ▲ 40 , ▼ 4 ) |
| miRNAs | 73 ( ▲ 39 , ▼ 34 ) | 10 ( ▲ 3 , ▼ 7 ) |
| *Others | 300 ( ▲ 201 , ▼ 99 ) | 54 ( ▲ 52 , ▼ 2 ) |
| *e.g. Pseudogenes, TEC, snoRNA, miscRNA, etc | Upregulation ▲ Downregulation ▼ |  |

### 4.2. UC mouse colon and blood transcriptomics signature

We identified 3,061 and 623 SDE genes from the total RNA-Seq and 73 and 10 SDE miRNAs from the small RNA-Seq data comparing UC mice colon and blood with control, respectively (Table 1). While SDE PCGs are mainly upregulated in both tissues, lncRNAs in colon are equally up- and downregulated and the miRNAs in blood are mostly downregulated. The RNA expression profile of the SDE genes and miRNAs, obtained by hierarchical clustering, is shown in heatmaps for colon and blood (Figure 1A, C). The SDE genes and miRNAs are highlighted in the volcano plots for colon and blood (Figure 1B, D).

**Figure 1:**
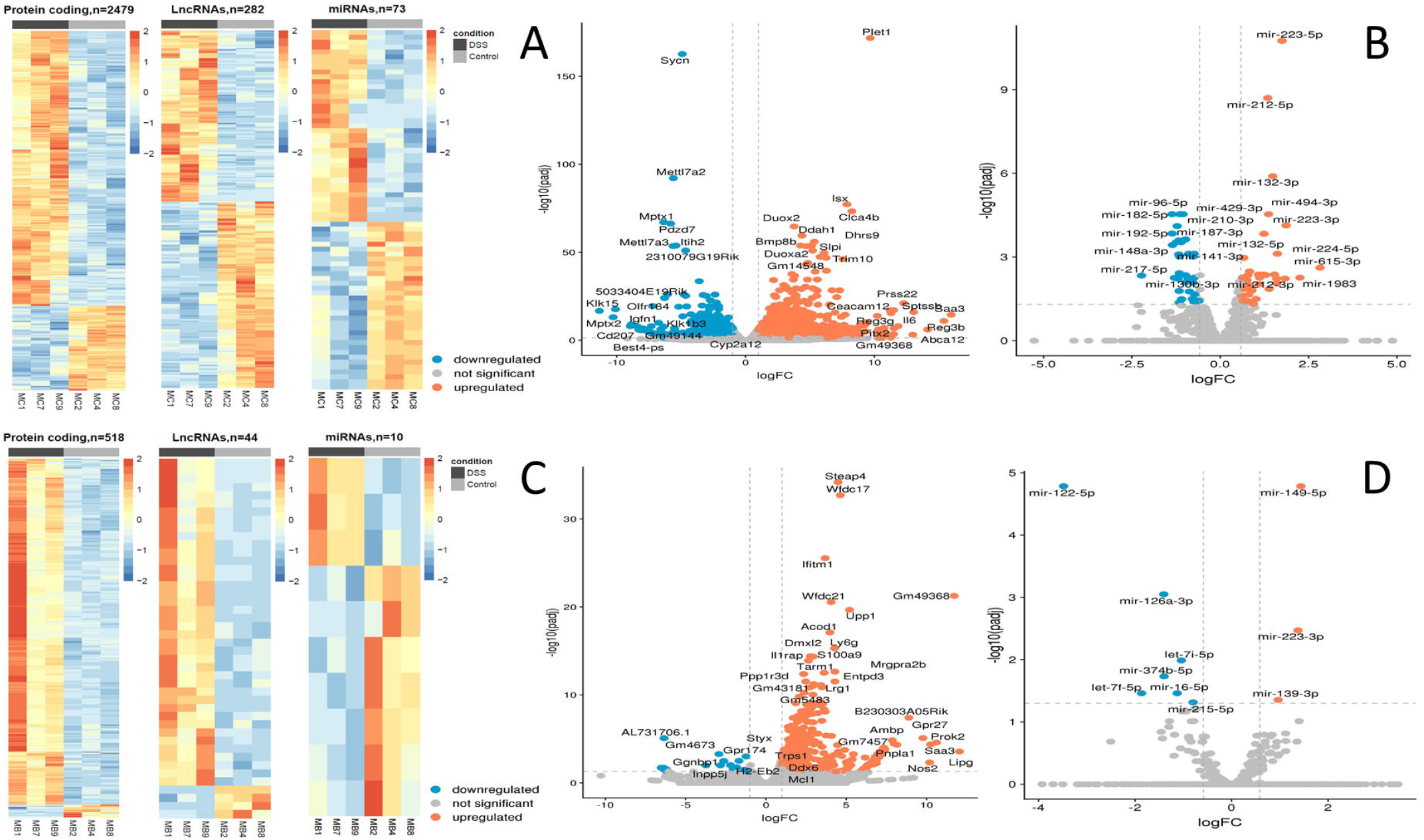
Heatmaps of SDE PCGs, lncRNAs and miRNAs, based on z-scores of normalized log counts for (A) colon and (B) blood. Volcano plots −log_2_FC ratio on the x axis versus −log10 padj on the y axis for colon (C) and blood (D).

From the SDE genes in colon, we detected 116 genes which have a human ortholog that is situated within 100 kb distance of the known Informatory bowel disease (IBD)-risk loci,^25, 26^ i.e. 110 PCGs, 2 lncRNAs and 4 other RNAs. In blood, 30 SDE genes, all PCGs, were IBD-risk loci associated (Supplementary File 3). Several of the SDE genes including *Reg3b, Abca12, Sptssb, Prss22, Pitx2, Ceacam12, Myot, Defb37, Gml* and *Ugt2b5* in colon and *Gpr27, Gm7457, Usp2, Stfa3, Gm37800, Dync2li1, Gm32486, Gm7206, Gm33326* and *Epha2* in blood have not been identified previously in this model. Top 10 up- and downregulated (padj≤0.05, avg. exp > 100) genes in colon and blood are listed in (Supplementary Table 1).

To investigate the roles of the SDE PCGs in UC colon and blood, we performed functional enrichment analysis (supplementary File 4). For mouse colon, the top ranked processes and pathways include immune and inflammatory system, response to inflammation and organ/tissue remodeling (Supplementary Figure 3A). In particular, the KEGG disease pathway for IBD is enriched with almost half of the pathway genes being SDE in our data. Although the number of SDE genes in blood is lower than in colon, the top ranked processes and pathways enriched for the SDE genes in blood overlap to a great extent with the colon (Supplementary Figure 3B) and are mainly related to the immune system, inflammatory system and connective tissue.

For the SDE lncRNAs, we performed functional enrichment of their genomic neighbors expressed in our samples (Supplementary Methods). We retrieved a total of 888 PCGs in colon and 159 PCGs in blood (Supplementary File 5). Cancer, organismal injury and abnormalities, gastrointestinal disease and immunological disease were overrepresented for both tissues with stronger significance in colon than in blood (Supplementary File 5).

In colon for 59 out of 73 SDE miRNAs, we identified 2,330 target candidates expressed in our samples and for blood, for 8 out of 10 SDE miRNAs we identified 1,641 targets expressed in our samples (Supplementary Figure 4A, B). Functional enrichment of the targets in colon and blood showed overrepresentation of immune related processes, several infectious and immunological diseases as well as JAK-STAT, thyroid hormone and MAPK signaling pathways (Supplementary File 6).

### 4.3. Overlap of SDE genes in colon and blood of the UC mouse

To identify genes commonly differentially regulated in colon and blood of UC mice with importance in the disease phenotype by overlap, 284 genes were retrieved (Supplementary File 7). Venn diagrams illustrating these genes’ relationship in colon and blood and the MA plot are shown in Figure 2A, B. From SDE common PCGs, *Prok2* (chemoattractant, labeled as outlier), *Saa3* (acute phase apolipoproteins) and *Gm49368* (predicted gene) were three most upregulated genes in the colon (logFC), while in blood, *Gm49368* was the highest followed by *Prok2* and *Saa3*. All 5 common lncRNAs showed significant upregulation with *Gm11714* most upregulated in blood and *Mirt2* most upregulated in colon (and second-most upregulated in the blood). For the miRNAs, mir-223-3p is the only common SDE (padj≤0.05) showing upregulation in both colon and blood with higher expression in blood than in colon, possibly indicating it is an immune-cell enriched miRNA. However, with a less stringent threshold of padj<0.1, we identify three other common miRNAs, downregulated in both colon and blood: mir-194-5p, mir-196b-5p and mir-215-5p. All three are highly abundant in colon and less abundant in blood and have previously been shown to be colon tissue specific in humans according to the miRNA TissueAtlas database.^27^ mir-223-3p showed to be more abundant in blood compare to the others.^27^

**Figure 2:**
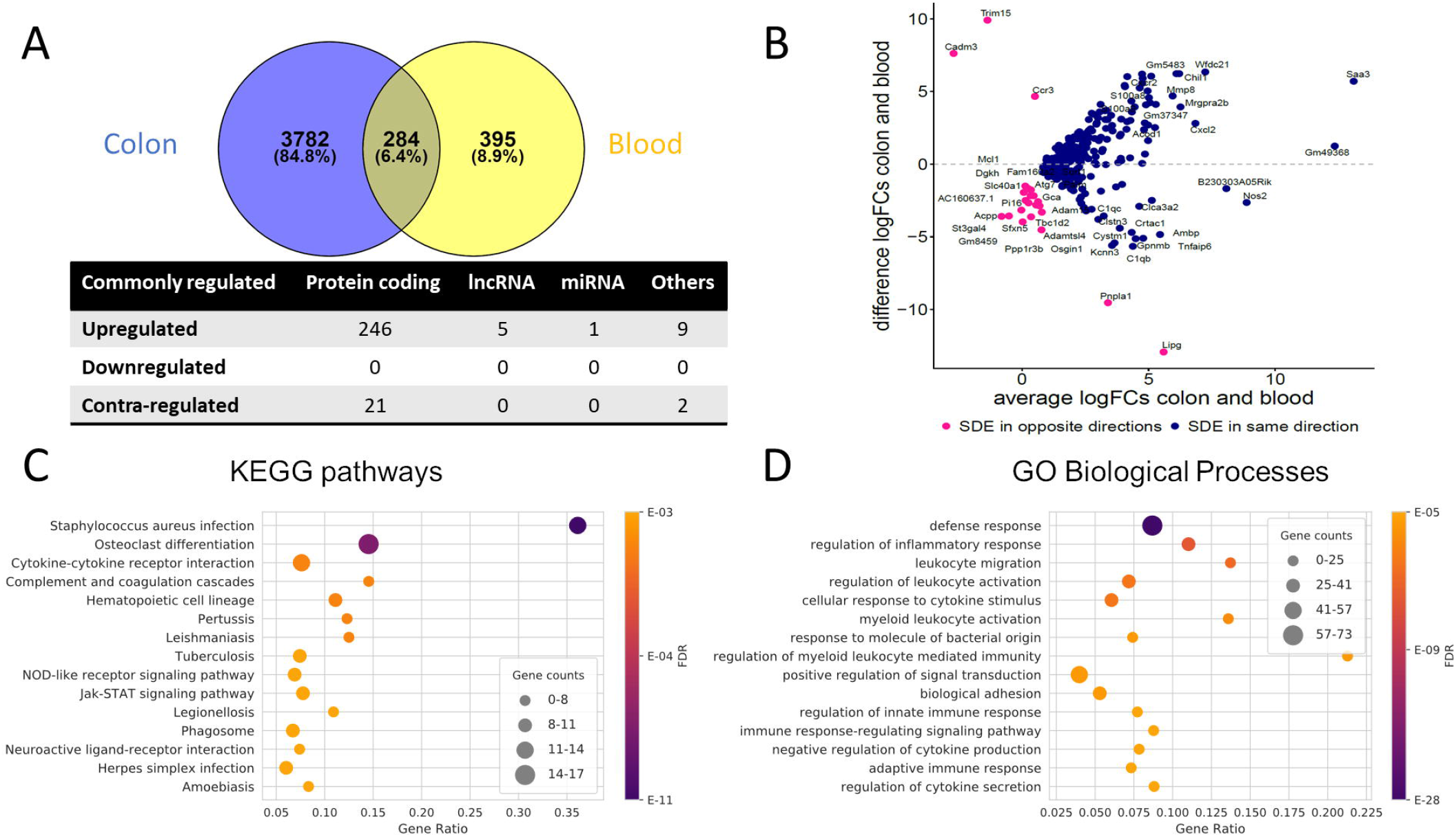
(A) Venn diagram of the SDE genes overlap between colon and blood; Other RNAs include pseudogenes, TEC, snoRNA, miscRNA, etc. Contra-regulated genes are commonly SDE genes in both tissues, which are regulated in opposite directions (B) MA Plot showing the relationships between log_2_FC in colon and blood for each of the common SDE genes; y axis indicates whether the gene has a higher FC in colon or blood, x axis reflects the cumulative FCs magnitude in colon and blood. Genes labeled as outlier in each organ were removed from the plot. (C) Top 15 enriched KEGG pathways and (D) Top 15 enriched GO biological processes for the common SDE gene between mouse blood and colon. Gene ratio corresponds to the ratio between the number of common SDE and all expressed genes in the pathway. The size of the dot indicates the number of SDE genes (gene counts), while the color represents the enrichment significance as given by the FDR adjusted p-values.

Collectively, the common SDE genes define distinct inflammatory, immune system and connective tissue related gene expression signatures. We observe enrichment of genes in processes related to the immune system, specifically regulation of inflammatory response and leukocyte activation, cytokine-cytokine receptor interactions (Figure 2C, D, Supplementary Figure 3C). Several KEGG pathways are related to infectious diseases (bacterial, viral and parasitic) that alter the immune response of cells. Analysis using IPA confirms these findings (Supplementary File 7).

### 4.4. Comparison between SDE genes in UC mouse colon and blood with human UC

#### 4.4.1. Colon and blood transcriptomics signature of human UC

In order to obtain reliable sets of SDE genes in human UC and to compare them with mouse in both colon and blood, we identified integrated sets of SDE genes using several public human datasets. For colon, 2 RNA-Seq^18^ and 2 microarray studies^19, 20^ and for blood, 2 RNA-Seq studies^16, 17^ were used (Table 2, Supplementary File 8). Human miRNA data for colon was obtained by combining one RNA-Seq and one microarray dataset. We have not been able to identify any high-quality miRNA dataset corresponding to whole blood of UC patients.

**Table 2:**
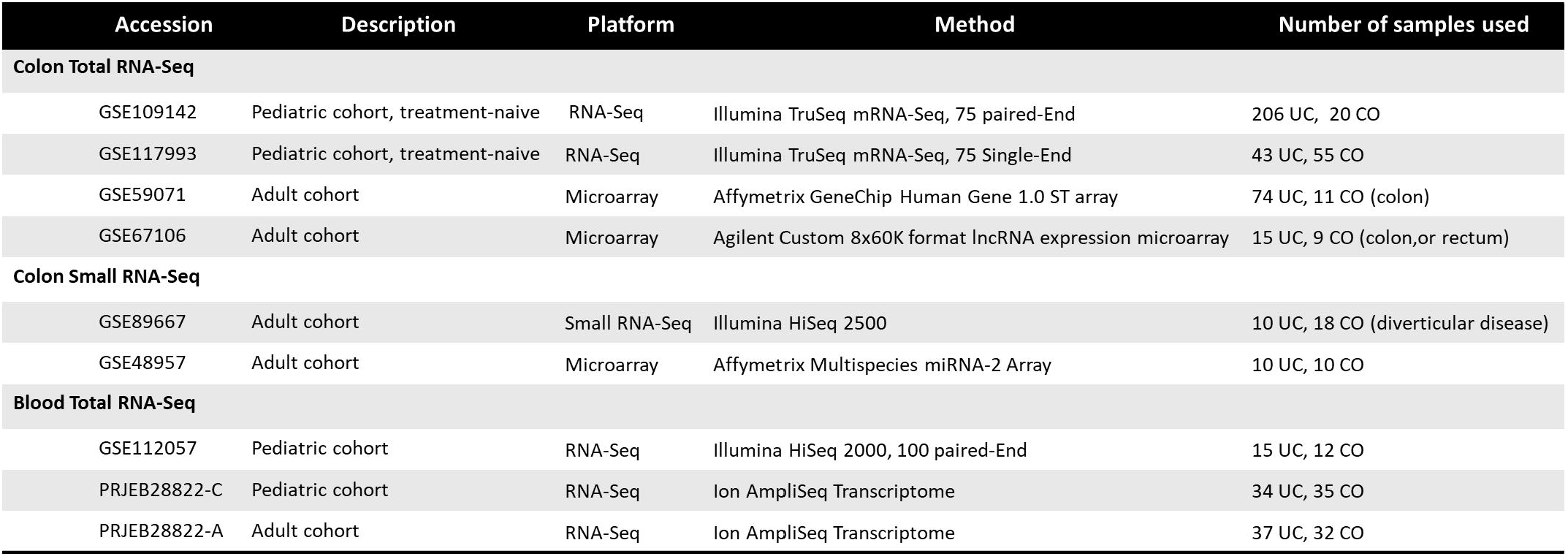
Public human UC datasets used in our study.

The combined sets of human SDE PCGs and lncRNAs, contained ~11,000 genes in colon, and ~2,000 genes in blood (padj≤0.05 in at least 2 datasets and consistently differentially regulated across datasets). We observed approximately equal proportions of up- and downregulated PCGs in colon, and a predominant (~75%) upregulation in blood. In both colon and blood lncRNAs were mainly (~60%) downregulated. Furthermore, ~2,500 genes in colon and ~130 genes in blood were inconsistently differentially regulated between the datasets and thus not included in the combined SDE gene sets. By combining the two public small RNA-Seq datasets in colon,^15, 21^ we obtained 207 SDE miRNAs with padj≤0.05 in at least one dataset and 37 were SDE in both datasets. No miRNA was found to be inconsistently differentially regulated between the two datasets (Table 3).

**Table 3:**
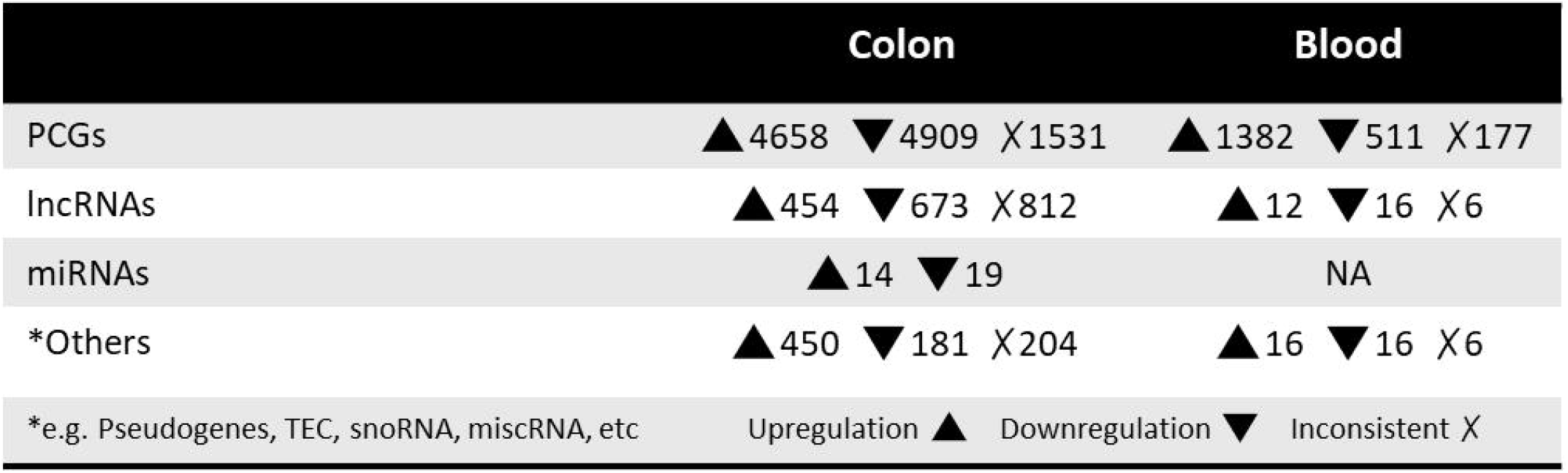
Numbers of SDE genes in human colon and blood combined datasets. (For all padj≤0.05 in min 2 datasets).

#### 4.4.2. Comparison between SDE genes of human and mouse UC

Using one-to-one and one-to-many orthology assignments, we compared the combined sets of SDE genes in humans with the sets of SDE genes in mouse colon and blood (Table 4). For colon, almost 75% of the SDE mouse genes with a human ortholog are also found in the human SDE set and for blood, only 30% of the SDE mouse genes with human orthologs are found in the human blood SDE gene set. The enrichment analysis for these common SDE PCGs in UC colon and blood showed mainly immune, inflammatory and connective tissue processes (Supplementary File 9 and 10). Seven lncRNAs showed significant up- (*H19, DNM3OS, 5430425K12RIK, 3110039I08RIK* and *FENDRR*) and down-regulation (*HOXA11OS* and *HOTTIP*) in colon of both mouse and humans, while no common lncRNAs were detected in blood. For 5 out of 7 common colon SDE lncRNAs, 33 neighboring genes (100kb upstream and downstream) were retrieved and they were enriched for terms including connective tissue disorders, organismal injury and abnormalities. From the common miRNAs, 12 were downregulated and 9 were upregulated in the same direction in the colon of mouse and human. Functional enrichment analysis for the 1,468 targets (detected in our samples) of 17 out of 21 SDE miRNAs in colon showed enrichment for terms including cancer, organismal injury and abnormalities, gastrointestinal disease, and several immune related processes (Supplementary File 9).

**Table 4:**
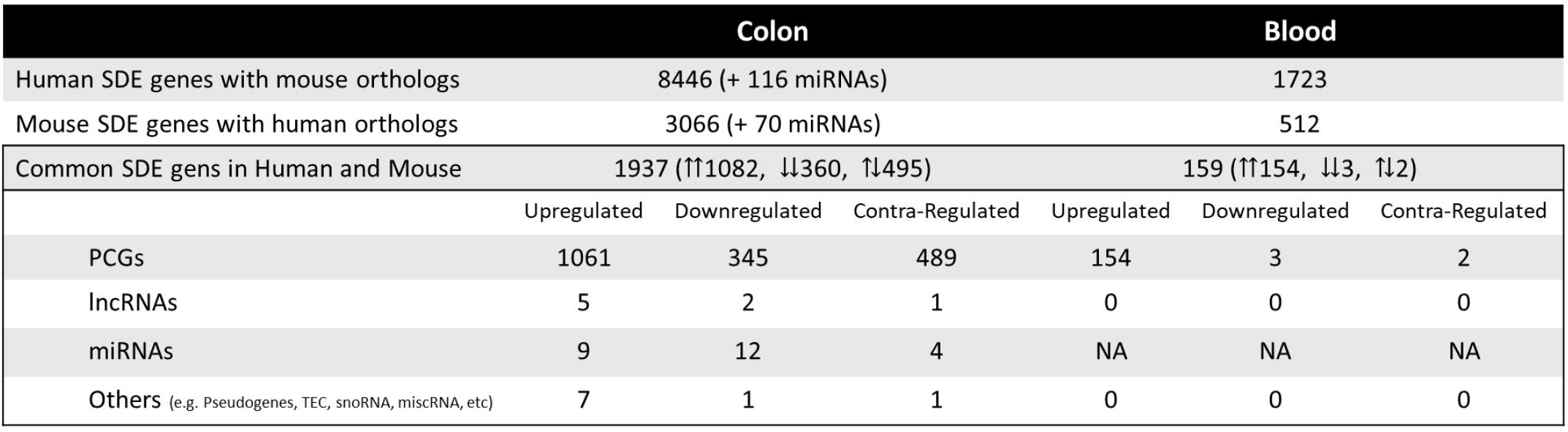
Numbers of overlapping genes between UC mouse and UC human colon and blood (For all padj≤0.05). ⇈: Upregulation in both mouse and human, ⇊: Downregulation in both mouse and human, ⇅: Opposite directions between mouse and human.

Though ~25% of common SDE genes in colon are contra-regulated, 356 genes with upregulation in mice and downregulation in humans showed to be mainly involved in lipid metabolism, ion transport, regulation of localization, trans-synaptic signaling. This could reflect species-specific differences in biological processes’ gene regulation. No enrichment was found for the 133 genes which were downregulated in mice and upregulated in humans. *Dio3os* was the only lncRNA in colon with upregulation in UC mouse and downregulation in humans. mir-10a-5p, mir-10b-3p and mir-10b-5p were colon miRNAs upregulated in mice and downregulated in humans, and highly expressed in both, while mir-130-3p was downregulated in mice and upregulated in humans, though lower expressed in both organisms compared to the other three miRNAs (Supplementary File 9).

Overall, 51 genes were identified as being commonly SDE between the colon and blood of both human and mouse when compared with the healthy controls (Figure 3A). These genes were all upregulated except for *PP1R3B* which is upregulated in blood and downregulated in colon. From these genes, only *SLC11A1* and *STAT3* were IBD risk loci associated. Collectively, these strictly filtered genes showed to be mainly involved in inflammatory, immunological and connective tissue related processes (Figure 3B, Supplementary File 11). Neither of 5 lncRNAs and 1 miRNA, which were commonly SDE between colon and blood of mouse, were present in the human datasets.

**Figure 3:**
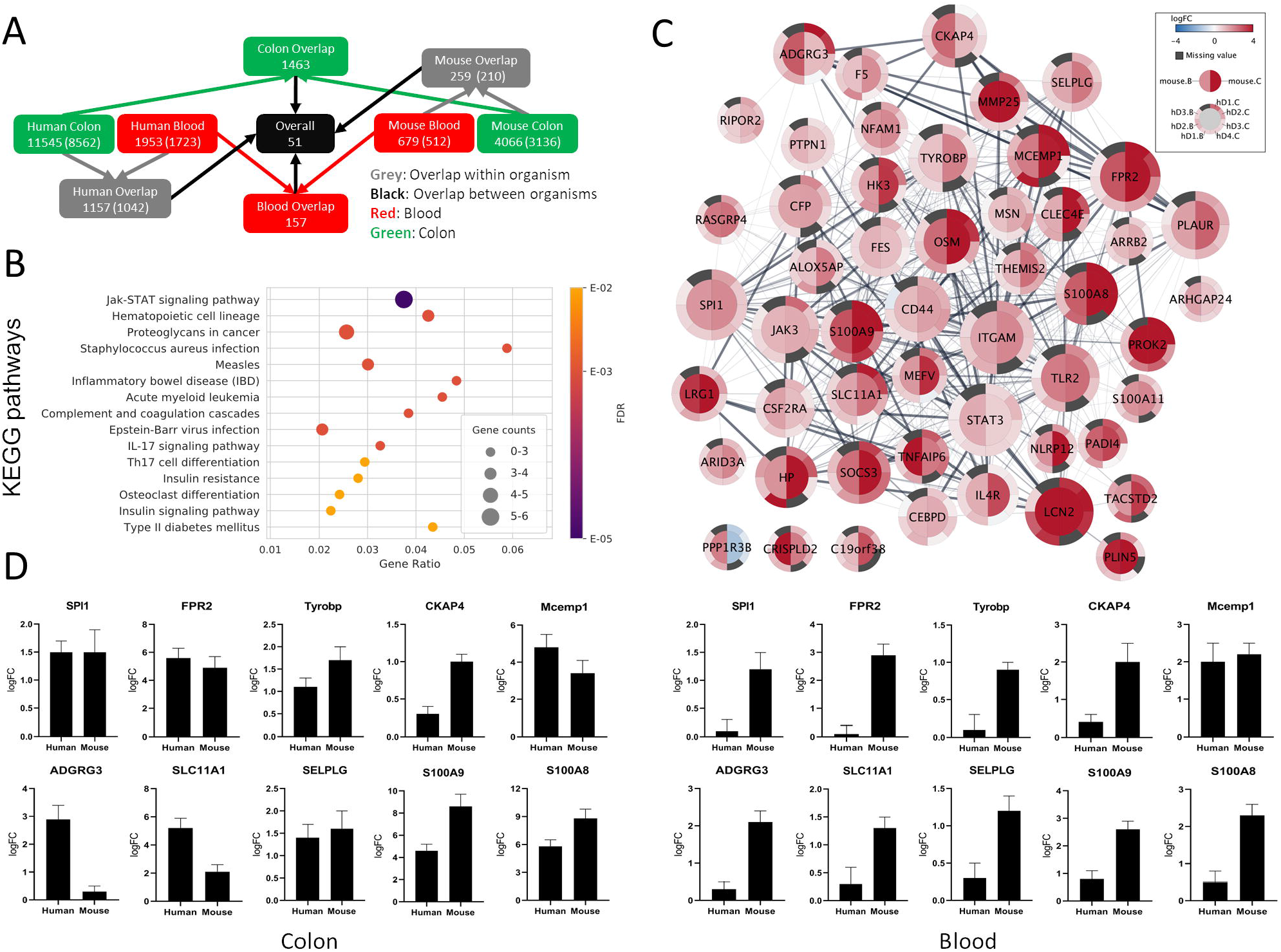
(A) Gene overlap diagram between human combined sets and mouse colon and blood SDE genes with padj≤0.05. Both the total number of genes in the sets and orthologous genes (in parenthesis) out of total is shown. (B) Top 15 enriched KEGG pathways (top 15 enriched GO biological processes) for the common SDE genes between human, mouse colon and blood. Gene ratio corresponds to the ratio between the number of common SDE genes in mouse-human and all human genes in the pathway. The size of the dot indicates the SDE genes number (gene counts), while the color represents the enrichment significance as given by the FDR adjusted p-values. (C) Differential gene expression visualization on the STRING network of genes common between mouse (inner node circle) and human (outer node circle) in blood (left node half) and colon (right node half). log2FC values are shown with a blue-white-red gradient and dark grey color indicates missing values. The size of the nodes corresponds to their network importance as measured by a combination of degree, closeness and betweenness centrality. (D) To validate the expression level of a few selected genes common in the colon and blood of human and mouse UC, qRT-PCR was performed. S100A8 and S100A9 were used as positive inflammatory controls.

From the 51 genes common between human and mouse in both colon and blood, 19 were selected for qPCR validation in a separate human UC cohort and the UC mice. Among these selected genes, 8 were genes that were either novel in a UC phenotype setting or not being previously reported as therapeutic target or biomarkers. Furthermore, they were central in the network of protein-protein associations retrieved from STRING v11 (Figure 3C, Supplementary File 11) as measured by a combination of three complementary centrality measures (degree, closeness and betweenness). While degree accounts for the direct interaction partners of each node, closeness and betweenness measure how central and thus important a node is with respect to all other nodes in the network (Supplementary Methods). The expression pattern of all tested genes by qPCR in colon and blood in both mouse and human validation cohorts nicely matched the RNA-seq analysis (Figure 3D and Supplementary Figure 5). The UC phenotype functional conservation and the importance for direct protein interactions with well-known immune-related and inflammatory genes suggests that these genes have high therapeutic-target and diagnostic application potential.

## 5. Discussion

Over the last few years, several transcriptional profiling studies have provided evidence regarding differential regulation of PCGs, lncRNAs and miRNAs in UC.^18, 20, 28, 29^ Given the complex nature of UC, an in vivo model approach to study its etiology is of paramount importance. As a surrogate to human UC, the DSS-UC mouse has been extensively used. However, its transcriptional landscape and its (dis-)similarity to human UC have not yet been studied and comprehensively characterized.

Here, we performed colon and blood transcriptional profiling of UC mice for both coding and non-coding RNAs and investigated the affected biological pathways. Our analyses showed that most of the differentially regulated genes are upregulated and involved in immunological and inflammatory responses as well as connective tissue, organ abnormality and injuries. This clearly points to the pathological nature of UC and importance of inflammatory processes in the destruction of epithelium and aggravation of disease. Moreover, alterations and mainly upregulation of genes which suppress inflammation, both in the colon and blood were detected. Genes, which demonstrated significant downregulation, were mainly genes with basic metabolic functions involved in cellular growth and proliferation, lipid, vitamin, amino acid and mineral metabolism. Most of the genes common between colon and blood of UC mice are regulated in the same direction (upregulated) and are related to immune or inflammatory processes. Among these genes, *S100a8, S100a9, Il1b, Lcn2, C3, Cxcl2, Nos2, Slpi, Socs3, Ifitm6, Csf2rb, Tlr2* and *Cd44* have previously been detected in UC mouse models in other studies.^30–34^ Several other SDE genes in colon and blood from our study including *Lipg, Gm5483, B230303A05Rik, Ambp* and *Trem3* have not been shown previously in this model. The largest overlap of 1,566 SDE genes was with a recent study by Czarnewski et al, where deep RNA sequencing^35^ was performed to stratify different subtypes of human UC using the common genes in human and mouse UC. Although they used a very similar approach, neither non-coding RNAs nor blood transcriptome were considered in their study.

As the first study investigating the transcriptional landscape of both lncRNAs and miRNAs in the colon and blood of the DSS-UC mice model, we identified widespread differential regulation of non-coding RNAs. Numerous SDE lncRNAs and miRNAs were identified in both tissues, many of which have not been reported previously. Among SDE lncRNAs in the colon, several are well-known, including *H19, Meg3, Hottip, Hoxa11os* and *Mirt2*. The latter was also SDE in blood and has been shown to be a negative regulator of inflammation.^36^ *H19* upregulation is believed to significantly decrease the expression of the vitamin D receptor and thus have a destructive effect on the intestinal epithelial barrier function by increasing permeability and decreasing the expression of ZO-1 and occludin tight junction proteins.^28, 37^ Not much is known regarding the other lncRNAs in UC settings, but recently *Meg3* was shown to inhibit the inflammatory response in ankylosing spondylitis characterized by chronic inflammation.^38^ *Hottip* silencing in rheumatoid arthritis led to reduced inflammation.^39^ In UC mouse blood lncRNAs *Neat1* and *Panct2* also showed to be SDE in disease. *Neat1* is shown to be pro-inflammatory and its inhibition suppresses the inflammatory response in IBD,^40^ whereas little is known about *Panct2*. Moreover, numerous SDE lncRNAs not reported before in UC mouse including top three upregulated *9130221F21Rik*, *E230034D01Rik*, *1200007C13Rik* in colon and *Gm7457*, *Gm37800*, *Gm32486* in blood and many more up- and downregulated new lncRNAs were identified in our study.

Among the SDE miRNAs, IBD-related miRNAs including miR-30, -223, -21, -142 were upregulated and miR-192 was downregulated in colon, while miR-223, -16 and -126 were upregulated in blood. miR-223-3p is the only miR upregulated in both colon and blood of the UC mice. It is upregulated in neutrophils and monocytes and acts as a controller of NLRP3 inflammasome activity, which regulates the intestine inflammatory process by affecting IL-1β production.^41^ It has also been shown that miR-223 mediates the cross talk between intestinal barrier and IL-23 pathway by targeting CLDN8, which is a claudin protein that constitutes the backbone of the intestinal barrier.^42^ miR-223 has also been used as a biomarker in IBD.^43^ Thus, evidence clearly suggests its proinflammatory role and highlights its potential as a RNA biomarker that seems to be conserved between different species. It is noteworthy that in the current study, many new SDE miRNAs not shown in this UC mouse model previously including top three upregulated mir-615-3p, mir-212-3p, mir-224-5p in colon and mir-149-5p in blood and many more are identified, which need future detailed functional investigations.

By comparing our UC mouse to human UC data, we identified similar widespread directional differential regulation of genes. The overlap of a substantial number of differentially regulated genes in mouse and human UC suggests fundamental common mechanisms responsible for disease onset and progression. To date, few studies using colon have compared the transcriptional changes in UC mouse models and human UC.^33–35^ *Holgersen et al.* identified 92 SDE genes in both human CD and UC,^33^ which were used for comparison to their three mouse models. In the DSS model, since colon samples were pooled, no statistical significance of differences could be calculated between disease and healthy mice. However, 59 genes showed to be common between our lists. In their study, non-coding RNAs and blood transcriptome were not studied. *Rankin et al*., however, identified 12 PCGs and 6 lncRNAs associated with DSS mouse colitis and human UC.^34^ No small RNA and blood profiling were performed in their study. In the *Czarnewski et al* study, 650 SDE genes common between human and mouse UC colon were identified, but no non-coding RNA and blood profiling was performed.^35^ Apart from these studies, to the best of our knowledge, there are no human-mouse UC comparative studies considering both colon and blood as well as coding and non-coding RNAs to the scale of our study.

Although we recognize that separate analysis of common genes in colon and blood may provide additional information, here, we focus on the genes that were common between both organs in the mouse and human and thus, may represent universal markers of UC. Overall, 51 common PCGs in mouse-human colon and blood with strong inflammatory and immunological profile showed to be consistent. Several of these genes, including *ITGAM, STAT3, LCN2, TLR2, PLAUR, JAK3, CD44, OSM, SOCS3, S100A8, S100A9, HP, CFP, IL4R, FES* and *CSF2RA*, have previously been associated to IBD. Interestingly, several less known/studied genes in IBD, including *SPI1, FPR2, TYROBP, CKAP4, MCEMP1, ADGRG3, SLC11A1* and *SELPLG* were identified among top candidates. *SPI1* encodes an ETS-domain transcription factor PU.1 that activates gene expression exclusively in hematopoietic cells including both myeloid and lymphoid cells e.g. during lymphocyte B cell development and is involved in disease like Inflammatory diarrhea, primary mediastinal B cell Lymphoma and pediatric T cell acute lymphoblastic leukemia.^44, 45^ *FPR2* works as a chemoattractant receptor involved in antibacterial host defense and inflammation through sensing of bacteria^46^ which is expressed not only by immune cells, but also epithelial, endothelial and fibroblasts cells^47^ that elicits proinflammatory responses. *FPR2* also promotes monocytes inflammatory activities and its absence in knock-out mice results in increased bacteria load in the liver and reduced neutrophil infiltration.^48^ *TYROBP* encodes a transmembrane signaling polypeptide associated with the killer-cell inhibitory receptor family and plays a role in signal transduction and inflammation.^49^ Loss of *TYROBP* has been shown to result in a presenile dementia with bone cysts.^50^ *TYROBP* expression is increased in Alzheimer’s disease (AD) patients and its deficiency in AD mouse showed to be neuroprotective and immune-inflammatory therapeutic which eventually slowed/arrested the progression of pathological late onset sporadic AD.^51^ *CKAP4* (also known as *CLIMP-63*) encodes a transmembrane protein and has been the focus of several investigations (reviewed in^52^) and is believed to regulate cell migration.^53^ *MCEMP1* encodes a single-pass transmembrane protein, which is involved in immune responses through mast cell differentiation.^54^ *ADGRG3* shows to regulate antimicrobial activity of granulocytes^55^ and seems to be required for macrophages local inflammation development.^56^ *SLC11A1*, previously known as *NRAMP1*, is expressed exclusively in immune monocytes and phagocytes like macrophage.^57^ Mutation in these genes may have a role in susceptibility to infections and autoimmune diseases.^58^ *SLC11A1* product acidifies the phagosome^59^ and thereby kills the entrapped pathogens. *SELPLG*’s strong association with the immune system has been emphasized before.^60^ This gene codes for *PSGL-1* protein, which is a counter receptor for P-selectin that facilitates immune responses by promoting immune effector-cells trafficking into inflamed tissue. These genes are evolutionary conserved and have known interactions with already well-known IBD candidate genes. Thus, these candidates potentially represent new UC diagnostic and therapeutic targets and novel avenues to more detailed disease mechanistic studies.

No long non-coding RNAs made it to the 51 final gene list. The major reason is that only a few lncRNAs are assigned orthology relationships across species in general (e.g. only 131 of all mouse lncRNAs are assigned a human ortholog based on HCOP database). However, we have been able to identify 7 lncRNAs that were SDE in colon in both humans and mice. Apart from H19 among them, not much is known about the other six lncRNAs in IBD, which clearly shows a need for detailed analysis. We also identified 21 miRNAs SDE in colon in humans and mice. Among them miR-146a-5p, -155-5p, 192-5p, 194-5p, 196b-5p, 200c-3p, 223-3p and 223-5p were frequently shown to be differentially regulated in IBD previously, which makes them promising candidate for further analysis.

Given the complexity of the whole colon and blood cell types, for distinguishing pathogenic mechanisms with higher resolution, one way could be to isolate different cell populations and perform single cell sequencing. One of the limitations of our study is that whole colon and blood gene expression profiling were performed. Albeit, the main rationale was to preserve the natural state of the disease as much as possible. In addition, the disadvantages for isolation of different cell types are the technical fractionation procedures and the time duration from tissue collection to sample processing, which potentially affect gene expression statuses specifically for non-coding RNAs in cells.

## 6. Conclusion

High-throughput transcriptome analysis provides a unique tool to discover new targets for therapeutic and diagnostic applications through identifying relationships between gene expression and disease phenotype. Here, we showed that a one-to-one comparison of the transcriptome of the DSS-UC mouse model to human UC can provide novel disease pathophysiology information by identifying genes and pathways with important contributions in UC. Our data suggest that prospective therapeutic interventions and diagnostic applications should target multiple major gene regulators involved in UC pathogenesis and propagations and combine several genes for valid biomarker applications. Moreover, targeting genes with conserved functional roles in the disease pathogenesis may offer a reliable UC treatment approach compared to targets with different functional roles in different tissues and organisms.

## Supporting information

Supplementary Section

## 8. Author contributions

R.Y. designed the study, established the mouse model, performed the experiments and interpreted the results and wrote the manuscript. (Conceptualization, Methodology, Software, Validation, Formal analysis, Investigation, Resources, Writing - Original Draft, Visualization, Project administration). O.P. processed and analyzed the RNA-Seq and public datasets and contributed to writing the manuscript (Software, Formal analysis, Data Curation, Writing - Original Draft, Visualization). N.T.D contributed to analyzing the data and to writing the manuscript (Software, Formal analysis, Writing - Original Draft, Visualization). C.A. provided support in analyzing the data and revised the manuscript. (Formal analysis). B.P. and U.H. provided tools and helps for establishing the mouse model. (Methodology, Resources). C.A.S.S performed the qPCRs. (Validation). A.H.M prepared the human samples and revised the manuscript. (Resources). T.L. provided essential insights and revised the manuscript. (Conceptualization). C.H.B.B and M.V prepared the human samples and revised the manuscript. (Resources). J.G. and L.J.J. provided essential tools and insights in designing the experiment, supervised the study and revised the manuscript. (Conceptualization, Supervision, Funding acquisition). F.P. contributed in designing the experiment, provided essential tools and insights, supervised the study and revised the manuscript. (Conceptualization, Supervision, Project administration, Funding acquisition). All authors read, reviewed and approved the final manuscript.

## 9. Acknowledgment

We would like to thank Dr. Charlotte Aaberg Poulsen from University of Southern Denmark for providing the animal study license for UC mouse model establishment. We also would like to thank Professor Poul Flemming Høilund-Carlsen and Christina Baun (Research radiographer) from Clinical Physiology and Nuclear Medicine at Odense University Hospital for performing microPET scanning of the UC mouse models. Moreover, special thanks to Giulia I. Corsi (Ph.D. candidate) from Center for non-coding RNA in Technology and Health for help with the visualization of enrichment analysis.

