## Supplementary Section for "Transcriptome profiling of ulcerative colitis mouse model suggests biomarkers and therapeutic targets for human colitis"

Reza Yarani[1*], Oana Palasca[2,3,4], Nadezhda T. Doncheva[2,3,4], Christian Anthon[3,4], Bartosz Pilecki[5], Cecilie A. S. Svane[1], Aashiq H. Mirza[3,6], Thomas Litman[7], Uffe Holmskov[5], Claus Heiner Bang-Berthelsen[8,9], Mogens Vilien[10], Lars J. Jensen[2,3], Jan Gorodkin[3,4], Flemming Pociot[1,3,11,12*]

[1] Translational Type 1 Diabetes Research, Department of Clinical Research, Steno Diabetes Center Copenhagen, Gentofte, Denmark
[2] Novo Nordisk Foundation Center for Protein Research, University of Copenhagen, Copenhagen, Denmark
[3] Center for non-coding RNA in Technology and Health, University of Copenhagen, Copenhagen, Denmark
[4] Department of Veterinary and Animal Sciences, University of Copenhagen, Copenhagen, Denmark
[5] Department of Cancer and Inflammation Research, Institute of Molecular Medicine, Faculty of Health Sciences, University of Southern Denmark, Odense, Denmark
[6] Department of Pharmacology, Weill Cornell Medicine, Cornell University, New York, NY 10065, USA
[7] Department of Immunology and Microbiology, University of Copenhagen, Copenhagen, Denmark
[8] Research Group for Microbial Biotechnology and Biorefining, National Food Institute, Technical University of Denmark, Kgs. Lyngby, Denmark
[9] Department of Gastroenterology, North Zealand Hillerød Hospital, Hillerød, Denmark
[10] Department of Surgery, North Zealand Hospital, Hillerød 3400, Denmark
[11] Copenhagen Diabetes Research Center, Department of Pediatrics, Herlev University Hospital, Herlev, Denmark
[12] Department of Clinical Medicine, Faculty of Health and Medical Sciences, University of Copenhagen, Copenhagen, Denmark

*All Correspondence to:

Reza Yarani:

Flemming Pociot:

### 1. Supplementary Materials & Methods

#### 1.1. The establishment and validation of the UC mouse model

Ten wild type (WT) C57BL/6 male mice (eight weeks old) were obtained from Taconic Biosciences (Ejby, Denmark). Mice were randomly group-housed under a controlled room temperature and 12:12 hours light–dark cycle with access to food and water ad libitum. The bedding was changed weekly. All experiments were carried out in accordance with and approved by the Danish Animal Experiments Inspectorate, Ministry of Food, Agriculture and Fisheries, Denmark at Odense University Hospital under 2015-15-0201-00730 license number.

Acute ulcerative colitis was induced using dextran sulfate sodium salt (DSS), colitis grade (mw: 36 kDa, CAS Number 9011-18-1, MP Biomedicals, Soho, OH, USA) by administration in the drinking water at a final concentration of 1.5% (wt/vol) for 7 days and then replaced by regular water. Control group was time-matched and received the same drinking water without DSS. If mice weight loss exceeded 20% of the initial weight, they were sacrificed by cervical dislocation. All animals were sacrificed by the same procedure on day 8.

Disease activity index (DIA) is the combined score for weight loss, stool consistency and bleeding, and the scores have historically correlated well with the pathological findings in the DSS-induced model of UC.[1] The

DAI in this study is based on the previously published scoring system,[2] which briefly is as follows. Weight: 0= no loss; 1= 5–10% loss; 2= 10–15% loss; 3= 15–20% loss; and 4 >20% loss; stool: 0= normal; 2= loose stool; and 4= diarrhea; and bleeding: 0= no blood; 2= presence of blood; and 4= gross blood. Blood in stool was assessed using the Hemoccult II test (Beckman coulter, Oakville, ON, Canada). The DAI scoring was performed from days 0 to 7 over the period of DSS treatment. For inflammatory response analysis customized ProcartaPlex immunoassays (Thermo Fisher Scientific) of inflammatory cytokines using Magpix multiplexing clinical diagnostics instruments for colon thickness culture and C-reactive protein for blood using Mouse C-Reactive Protein/CRP Quantikine ELISA Kit (R&D Systems, Cat: MCRP00) were measured.

#### 1.2. RNA extraction from blood and colon and quality control

Based on the DAI, six mice (3 UCs and 3 controls) were selected. Blood from the mice was collected in RNAprotect Animal Blood Tubes (Cat No./ID: 76544, Qiagen). Colons were collected, weighted and the lengths were measured and immediately preserved in RNAlater RNA stabilization Reagent (Cat No./ID: 76104, Qiagen) and stored at -80 C.

Total RNA (including miRNA) from blood was isolated using RNeasy Protect Animal Blood Kit (Cat No./ID: 73224, Qiagen) with Buffer RWT (cat. no. 1067933, Qiagen). Colon total RNA was isolated using miRNeasy Mini Kit (Cat No./ID: 217004, Qiagen) after simultaneous tissue disruption and homogenization using TissueLyser LT (Cat No./ID: 85600, Qiagen) and Stainless-Steel Beads, 5 mm (Cat No./ID: 69989, Qiagen). RNA quantity was measured using the NanoDrop 2000c spectrophotometer (Thermo Fisher Scientific™, Waltham, Massachusetts, USA). and RNA integrity (RIN quality) was checked by Agilent 2100 Bioanalyzer system for both total and small RNA using Agilent RNA 6000 Nano Kit (5067-1511, Agilent) and Agilent Small RNA kit (5067-1548, Agilent) for all samples.

#### 1.3. Library preparation, RNA sequencing

Library preparation for colon RNA samples was performed using Ribo-Zero™ Magnetic Gold rRNA Removal Kit (Epidemiology, MRZE724, illumina) and for blood, using Globin-Zero Gold rRNA Removal Kit (GZG1224, illumine) based on the standard kits protocol. Total and small RNA sequencing were performed by BGI, Hong Kong using the illumina HiSeq 4000 system default protocol of sequencing.

#### 1.4. Data processing and differential expression analysis

For the total RNA-Seq data, raw fastq files were cleaned for the specific Illumina TruSeq adapter content and trimmed for low quality tails using Flexbar[3]. Between 3-4% of total raw reads had been removed in each sample in the cleaning step, resulting in a total of 40M-66M paired-end reads per sample. Next, reads of ribosomal origin were removed using the bbduk software (with option k=31) from the bbmap library (https://sourceforge.net/projects/bbmap/), by matching reads to a rRNA collection of sequences extracted from the SILVA database[4], as described in https://www.biostars.org/p/159959/. Between 1%-29% reads of possible ribosomal origin were removed from each sample. The remaining reads were mapped to the mouse genome (GRCm38) using STAR version 2.5.2a.[5] (options: --outSAMstrandField intronMotif; --outSAMunmapped Within; --outSAMattrIHstart 0; --outFilterType BySJout; --outFilterIntronMotifs

RemoveNoncanonical; --alignSoftClipAtReferenceEnds No), yielding total mapping rates between 90%-97%, and uniquely mapping rates between 33%-88%, the latter corresponding to library sizes between 17M-58M reads. Transcript and gene level quantification was obtained using Stringtie (version 1.3.4c)[6] in estimate mode (-e option) with gtf annotation files corresponding to Ensembl (release 92)[7] and default options. Gene-level raw counts were obtained using the prepDE.py script provided together with Stringtie. Gene biotypes were extracted from the gtf annotation file corresponding to Ensembl (release 97).

For the small RNA samples, raw fastq files were trimmed with Flexbar[3] for the specific adapter sequence ("AGATCGGAAGAGCACACGTCT"), yielding between 15-25M cleaned reads per sample. The reads were further collapsed using fastx-toolkit and mature mouse miRNAs annotated in miRBAse version 22[8] were quantified using the quantifier.pl module from the miRDeep2 software, version 2.0.1.0[9] , default options. Between 14-20M reads per sample mapped to the set of mouse miRNAs annotated in miRBAse v.22.

For both total and small RNA-Seq, differential expression analysis was performed using the DESeq2 package version 1.22.2[10] ,using raw gene-level counts as input for total RNA-Seq and mature miRNA counts for small RNA-Seq. A simple control-treatment design was used, with Wald test for log2FC different from 0 and the default Benjamini-Hochberg FDR control method. The genes marked as weakly expressed by the independent filtering implementation in DESeq2 were not included in further analysis. The genes detected as outliers, when enabling outlier detection in DESeq2 (cooksCutoff=TRUE) were also not included, except for when the gene was expressed in at least two of the disease samples.

#### 1.5. Functional enrichment and network analysis of differentially expressed PCGs

Functional enrichment of KEGG[11], Reactome[12] and Gene Ontology (GO) annotations[13] was reported by the stringApp v1.5.1 in Cytoscape 3.7.2 for terms with an adjusted p-value < 0.05. A custom background consisting of all expressed genes in the respective tissue was used. The enrichment was determined using Fisher's exact test and the resulting p-values were corrected using Benjamini-Hochberg multiple testing correction[14]. Additionally, GO annotation terms with less significant adjusted p-value and more than 25% of gene overlap with a term with more significant adjusted p-value were considered redundant and filtered out.

Enrichment of diseases and disorders, molecular and cellular functions, and physiology system development and function annotations came from Ingenuity pathway analysis (IPA) software program v51963813 [15]. For IPA, terms were grouped in higher order categories and the range of adjusted p-values (Benjamini-Hochberg multiple testing correction) was reported with a threshold of padj < 0.05. Ingenuity knowledge base (gene only) was used as the background. Ingenuity knowledge base is a comprehensive database of genetic data manually curated from the literature and/or third-party databases.[15]

For the STRING networks of SDE in mouse colon, mouse blood as well as common between mouse colon and blood, the default interaction confidence score of 0.4 was used in stringApp v1.5.1 in Cytoscape 3.7.2. The resulting networks were clustered by the Markov Cluster Algorithm (MCL) using the clusterMaker v1.3.1 app[16]. The STRING combined confidence scores were used as weights for the clustering and the inflation factor was set to 4. The top 5 clusters (based on size) were selected and further analyzed for functionally enriched terms (using the network they belong to as background) among the annotation resources available in the stringApp enrichment with focus on GO Biological Process, KEGG and Reactome pathways. The Omics Visualizer app v1.3 [17] was used to show the logFC values in the network visualizations.

In order to rank the SDE genes common between human-mouse blood and colon based on their importance, we retrieved a STRING V11 network for all edges among them with a score confidence above 0.0. Then, we computed three different centrality measures (weighted degree, betweenness and closeness) for each node and aggregated the three rankings using Borda count to obtain a final overall rank for each gene. In particular, the weighted degree corresponds to the sum of edge scores for the edge incident to a node and can be used to identify genes whose gene products interact with many other proteins. Closeness centrality is computed as the average shortest path distance of a node to all other nodes in the network and is normalized by the number of nodes in the network. This measure can highlight genes that have an important role for the connectivity of the whole network. Betweenness centrality is a measure of the importance of a node for the shortest paths connecting other nodes in the network and can highlight nodes that have the role of bottlenecks. It is computed as the average number of shortest paths a node is part of compared to all shortest paths in the network. It is further normalized by the number of all node pairs in the network. For closeness and betweenness centrality, the edge scores were converted to distances using the transformation -log(score). To combine the three complementary centrality measures, we used Borda count[18], a positional voting system, which rewards the most points to the candidate with the highest rank. When all candidates have been ranked and given points for all measures, the points are added up and the candidate with the most points wins. The resulting ranking of the combined centrality measures indicates which genes are most central in the network and thus most important for the interactions between all genes in the network.

#### 1.6. Analysis of lncRNAs through guilt by association and functional enrichment

To Investigate the possible roles for SDE lncRNAs in the UC mouse through chromosomal proximity with annotated genes, the guilt by association principle was performed.[19, 20] We define for each tissue the sets of SDE lncRNAs as (|log2FC|>1, padj ≤0.05) and retrieved their genomic protein-coding neighbor genes within a span of 100 kb upstream and downstream, which were expressed in our dataset. In the functional enrichment analysis using the stringApp in Cytoscape, we used as background all expressed genes in the respective mouse tissue and significance threshold of adjusted p-values < 0.05. For the functional enrichment with IPA, we used the whole genome (ingenuity knowledge base, gene only) as the background and a significance threshold for adjusted p-values < 0.05.

#### 1.7. Identification and functional enrichment analysis of miRNA targets

A list of potential targets for the significantly differentially expressed (SDE) miRNAs ( padj≤0.05) was obtained by combining experimentally validated miRNA-target interactions from DIANA-TarBase v8 [21] and MirTarBase v8.0 [22]. For DIANA-TarBase, we selected the miRNA-target pairs that are supported by direct interaction assays (HITS-CLIP or luciferase reporter assay) and required at least two occurrences in the database for each pair coming from HITS-CLIP experiments. For MirTarBase, we only included the pairs labeled in the database as "strongly supported", which includes interactions identified by Western blot, qPCR or Luciferase or GFP reporter assays. This initial set of miRNA-target pairs contained information for 374 miRNAs, 5965 mRNAs and summed up to 36,970 pairs. After intersecting the initial set of miRNA-target pairs with our expression data, in colon, we obtained a set of 59 miRNAs, 2330 targets, and 5369 miRNA-target pairs. In blood, we obtained a set of 8 miRNAs, 1641 targets, and 2386 miRNA-target pairs. In the functional enrichment analysis performed with the stringApp, we used a custom background set of 11,631 genes consisting of the mRNAs occurring in at least one direct interaction in TarBase or occurring among the strongly supported interactions in MirTarBase.

MiRNA-target interactions for SDE miRNAs (padj<0.05) and their top targets were also visualized using Cytoscape for each of the tissues. For visualization purposes, we reduced the large set of putative targets for some of the miRNAs by showing a maximum of 5 targets for each miRNA. We chose the top 5 after sorting the targets by their average expression in the control samples. The rationale for using this measure was the assumption that miRNAs will tend to bind to the highest expressed of their targets. We estimated the abundance of the targets using geTMM units (gene length corrected trimmed mean of M-values).[23] For obtaining these units, we followed the instructions from Smid et al[23] and divided the raw gene counts (as obtained from StrigTIe prepDE.py script) by gene length, and then used these gene-length corrected counts in edgeR v.3.5 [24] to obtain TMM-values.

#### 1.8. Identification of IBD loci associated lncRNAs, miRNA and PCGs

We extracted a list of 245 unique IBD loci reported by de Lange[25] and Jostins[26] from the GWAS catalog[27] on March 1st 2019. For each locus, we retrieved PCGs, lncRNAs and miRNAs, within a span of 100 kb upstream and downstream of the strongest SNP signal, using the human gene set annotation file corresponding to Ensembl 97, resulting in 1,607 human genes. Out of these, 716 human genes have a mouse ortholog (one-to-one or one-to-many), which correspond to 725 mouse genes located within the 100kb distance of the risk loci. Finally, we checked which of the neighbors are expressed in our mouse data.

#### 1.9. Analysis of the human datasets

##### 1.9.1. RNA-Seq datasets

For the RNA-Seq datasets, we obtained the fastq files from the European Nucleotide Archive.[28] A preprocessing step of the data included adapter trimming (where not already trimmed), removal of reads mapping to a collection of ribosomal RNAs as described in section 1.4, as well as removal of the reads mapping to HBA1, HBA2 and HBB genes in blood samples. Genes were further quantified using Salmon v0.13.1 [29] in quasi mapping mode and setting the --validate Mappings parameter, using an index built from the cDNA sequence of coding and non-coding transcripts of Homo_sapiens.GRCh38.

Differential expression was performed using DESeq2, using the likelihood ratio test (lrt) hypothesis testing, with gender + condition as full model and gender as a reduced model. Except for dataset GSE109142, where gender was provided by the authors, we inferred the gender using the sex-specific genes RPS4Y1, EIF1AY, DDX3Y, KDM5D, and XIST, as indicated in.[30] Using principal component analysis, the expression profile of these genes produced two clearly distinct clusters on the first principal component, capturing 99% of the variance in the data.

Dataset PRJEB28822, contained both samples corresponding to pediatric patients and samples from adult patients, and we performed the differential expression separately for the two groups. After removing rRNA and hemoglobin genes, the library size of the samples had a wide variability range, from 0,4 million to 22 million mapped reads. Based on a balance we observed between the number of samples vs. a minimum coverage required for each sample, we opted for removing samples with less than 7,5 million mapped reads. We have also removed 18 samples with a mapping rate below 50%, and where the reads mapped exclusively in a forward manner, while the rest of the samples were not strand specific. After filtering, we ended up with 101 samples for adults (32 control, 32 CD and 37 UC) and 102 samples for children (35 control, 33 CD and 34 UC).

Some of the datasets contained samples from both Ulcerative Colitis and Crohn's disease patients. We used the CD samples as well in the normalization of the data, but the reported results are based on comparing UC vs. control samples.

For the small RNA-seq datasets, we obtained the raw fastq files studies from the SRA archive (see Table 2 for accession numbers) and applied the same pipeline as described in Section 1.4, using the GRC38 Gencode human genome. For trimming, we identified the specific adapters from each dataset.

##### 1.9.2. Microarray datasets

For the public microarray data analysis, we used the GeoQuery package (v.2.50.5)[31] to retrieve the already normalized intensities, and the Limma package (v.3.38.3)[32] for differential expression analysis, with a simple model design based on condition (UC, CD or Control).

In order to standardize gene identifiers, we mapped probesets to Ensembl gene IDs. Where two or more probesets mapped to the same gene ID, we selected the probeset with the highest average expression value. We did the filtering of duplicate probesets and control probe sets before generating the results tables, such as not to bias the p adjusted values. When the same probeset could map to several genes, we included all the genes in the result file.

For the dataset GSE59071, the probeset gene mapping corresponding to the Affymetrix Human Gene 1.0 ST Array was obtained from Ensembl Biomart, allowing therefore for a direct mapping between probeset and Ensembl gene IDs.

For the dataset GSE67106, we mapped the Gene.name column (containing either gene name or Ensembl transcript IDs) to Ensembl gene IDs, using dictionary files extracted from Ensembl Biomart. Where mapping was not possible, we tried mapping the RefSeq ID columns to the Ensembl Gene ID. For many of the patients included in this study, samples from multiple locations of the intestine were available. We selected only a subset of the samples, such that we included only one sample per patient, preferably from colon sigmoideum, but also colon transversum or rectum where colon sigmoideum was not available.

For small RNA microarray datasets, in order to synchronise the miRNA identifiers from the older microarray studies to the current miRBase v.22 identifiers, we mapped the mature miRNA sequences provided in the microarray annotations to the set of mature human miRNAs in miRBase, and retained only unique sequences for further analysis.

##### 1.9.3. Combining multiple human datasets

We obtained a set of human differentially expressed genes for each tissue (colon and blood) by generating a union of the differentially expressed (DE) genes from each public human dataset for this tissue. First, for the mRNA-Seq datasets, we obtained all DE genes with padj $< 0.05$ and a minimum average expression across samples of 2 counts in any of the datasets (here, counts = number of reads assigned to the gene, normalized with library size; the rationale for setting a threshold was to eliminate very low expressed genes). We kept track of the number of datasets, in which each gene was DE, and whether the gene was consistently differentially regulated (in the same direction) or inconsistently differentially regulated (up-regulated in one dataset and down-regulated in another dataset) across datasets. A set of combined SDE genes in blood and colon was finally extracted, requiring each gene to be SDE in at least two datasets; inconsistently differentially regulated genes were excluded. For the small RNA-Seq datasets, since we were only able to combine two datasets, we only required that each miRNA is SDE (padj$\leq$0.05) in at least one of the datasets.

#### 1.10. Comparison between mouse and human data

The final combined human lists were intersected with the mouse SDE blood and colon gene sets (padj<0.05) using orthology relationships extracted from Ensembl (release 97).[33] For one-to-many orthology relationships, we first selected the gene with the highest average expression across samples from its orthology group, irrespective of whether that gene was differentially expressed or not. For the human combined datasets, we used the average expression from only one of the datasets (the deepest sequencing RNA-Seq dataset - GSE112057 in blood and GSE109142 in colon) for selecting the highest expressed genes in each orthology group.

Due to the lack of orthology assignments for lncRNAs in Ensembl, we supplemented our orthology set with all the orthology predictions available for lncRNAs in the HCOP database.[34, 35] In total, 131 mouse lncRNAs with a one-to-one ortholog in humans, and 3 mouse lncRNAs with one-to-many or many-to-many orthologs were extracted. For miRNAs, we inferred correspondence between mouse and human miRNAs using the miRBAse mature miRNA name and intersected the human lists with the mouse SDE miRNAs (padj≤0.05).

#### 1.11. Sample collection for patients and controls

Colon biopsies and blood samples from UC and control individuals were collected from a cohort at North Zealand Hospital, Hillerød, Denmark (described here[36]). The study was approved by the Regional Ethical Committee (H-4-2012-030). Prior to the sample collection and medical history recording, written informed consent was acquired from all the participants. Overall patient demographics can be seen in Supplementary File 1. Only subjects with no signs of autoimmune or inflammatory disease were included as control individuals. In UC patients, one to five endoscopic colon (colon sigmoideum) and rectum pinch biopsies were extracted from the macroscopically inflamed mucosa. The biopsies inflammation status was confirmed by histologic examination (pathologists were blinded to the status of inflammation) and features of chronic intestinal inflammation for each patient were scored using a previously described scoring system for UC.[37] For the control group, biopsies were taken from the same locations. All biopsies were placed in the RNAlater solution (QIAGEN, Hilden, Germany), and stored at -80ºC until processing. Additionally, to further confirm the biopsies inflammation status, a panel of 26 pro-inflammatory markers (cytokines, interleukins, metalloproteases) using quantitative real-time PCR (qPCR; Fluidigm platform) was used (data not shown).

#### 1.12. Validation of SDE genes by quantitative real-time PCR

For quantitative real-time PCR (qRT-PCR) hydrolysis probe-based designed PrimeTime qPCR 5′ Nuclease assays procured from Integrated DNA Technologies and predesigned TaqMan gene expression assays from Thermofisher. The double-quenched hydrolysis probes with 5′ FAM fluorophore, a 3′ IBFQ quencher, and an internal ZEN™ quencher were used. All complementary DNAs (cDNAs) were prepared using high capacity RNA to the cDNA synthesis kit (Thermofisher, Cat: 4387406) following the manufacturer's instructions. qRT-PCR was performed with 5 ng of cDNA per well template with TaqMan™ Gene Expression Master Mix (Thermofisher, Cat: 4369016). For PCR amplification, the following thermal profile was used: 2 minutes at 50°C; 10 minutes at 95°C; 40 × (15 second at 95°C, 60 second at 60°C). The qRT-PCR results were statistically tested with a Student's t-test. A p-value <0.05 was considered statistically significant.

### 2. Supplementary Results

#### 2.1. UC Mouse model validation

On average, DSS-UC mice lost 13% of their weight, 28 mg of colon dry weight and 1.7 cm of colon length compared to the healthy age-matched group. Immunohistochemical staining of the distal colon demonstrated that in healthy controls, crypt architecture was intact, and no immune cell infiltration could be seen, while UC mice presented high levels of immune cell infiltration, with large amounts of mononuclear and granulocytic leukocytes and significant ulcer.

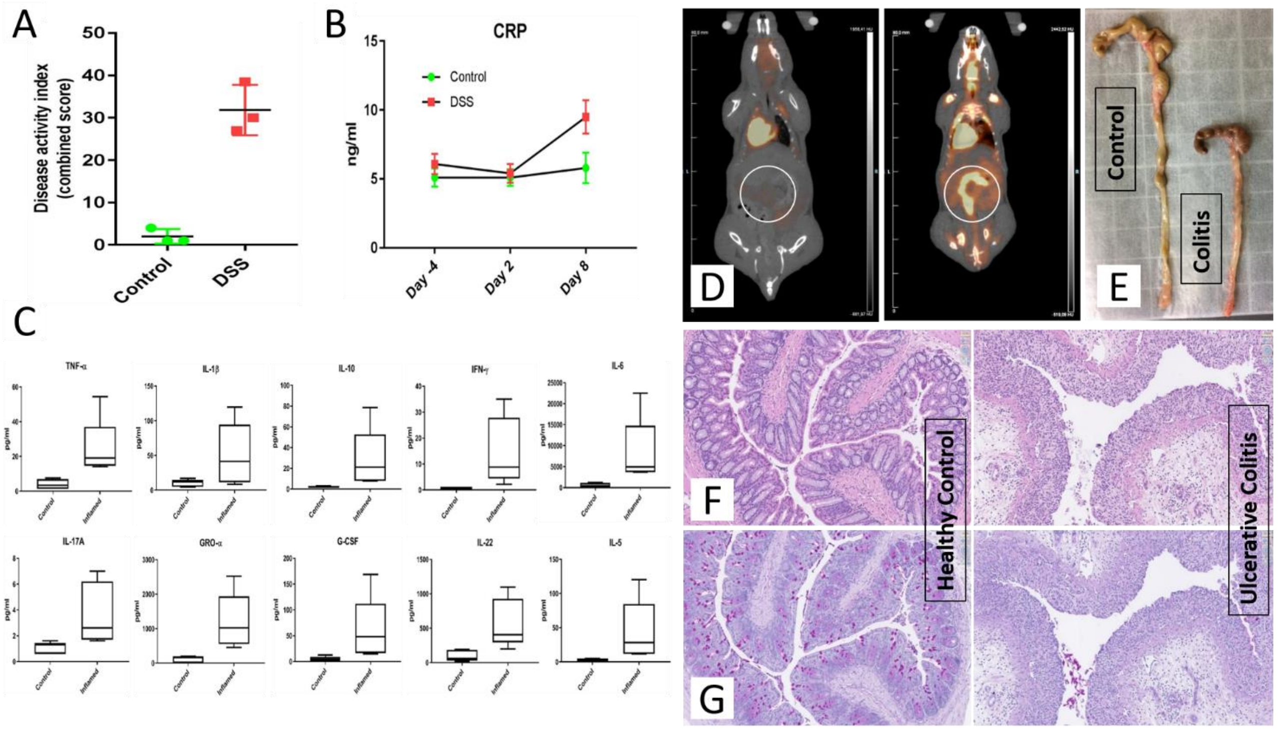


**Supplementary Figure 1.** UC Mouse model validation. (a) Disease activity index (DAI) score. Several clinical indexes of disease were monitored to calculate a cumulative DAI score. (b) peripheral blood C-reactive protein level measured for the control and DSS groups to assess the temporal inflammation induction (c) A set of Pro- and anti-inflammatory cytokines including TNF-α, IL-1β, IL-10, IFN-γ, IL-6, IL-17a, GRO-α, G-CSF, IL-22, IL-5 were measured by luminex assay for the model validation using colon tissue thickness culture (d) $^{18}$F-FDG microPET scanning was used to confirm the inflammation induction in the model. (e) Colon length difference between healthy controls and UC mice model. Histopathological changes including progressive superficial inflammation, epithelial and goblet cell erosion and immune cell infiltration was vastly observed in (f) H&E and (g) PAS stained colons of disease mice.

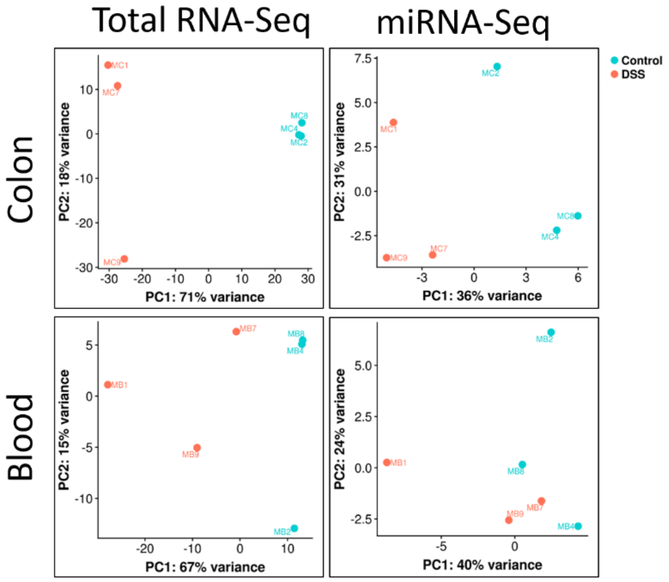


**Supplementary Figure 2.** PCA analysis based on the top 500 highest variance genes, using rlog transformed counts. Each data point is a separate mouse. In the total RNA-Seq colon and blood, as well as in the small RNA-Seq colon, control (n=3) and DSS (n=3) mice are clearly separated on the first principal component. The two groups are not well separated in small RNA-Seq blood on either of the first two components, in accordance with the lower number of SDE miRNAs between DSS and Control mice.

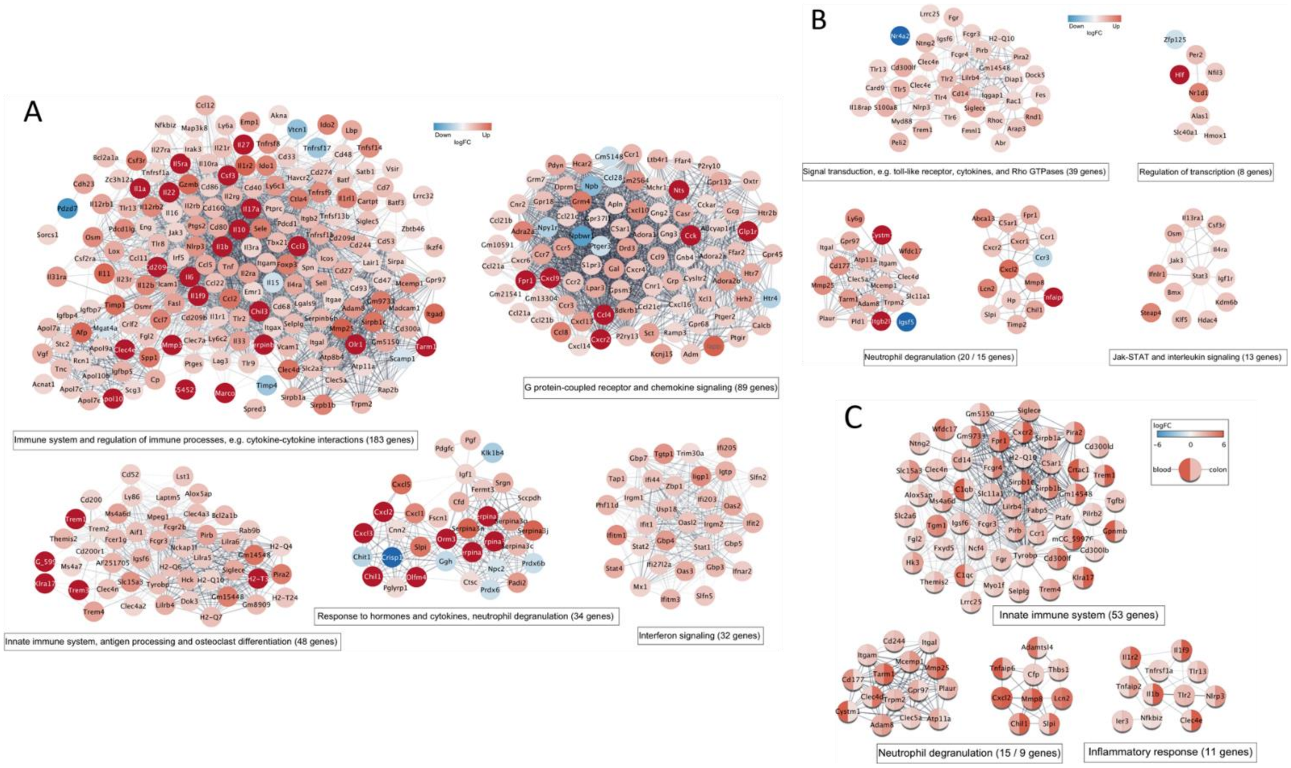


**Supplementary Figure 3.** Network visualization and functional enrichment of the SDE PCGs. The networks of functional associations between the SDE genes for (A) mouse colon, (B) mouse blood, and (C) the colon-blood common PGCs were constructed using the stringApp and visualized in Cytoscape. The logFC expression values in each tissue are shown as node colors using a blue-white-red gradient. For each network, the top 5 clusters (groups of strongly connected genes) were identified by Markov clustering (MCL) using the clusterMaker app and annotated based on their functional enrichment retrieved with stringApp. The resulting clusters represent different groups of genes involved in specific immune related processes.

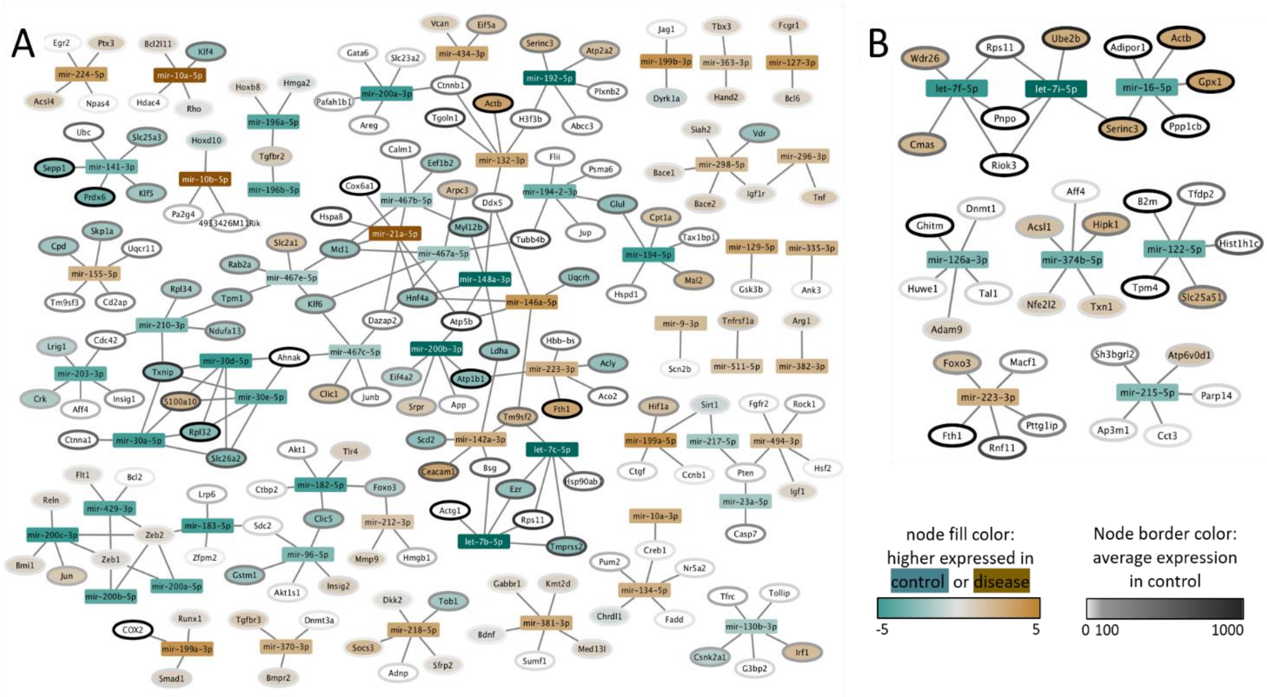


**Supplementary Figure 4.** Network visualization of miRNAs and their top 5 targets selected based on their average expression in control samples for (A) mouse colon and (B) mouse blood. miRNAs are shown as rectangles and PCG as oval-shaped nodes. The node border color indicates the average expression in the control samples. The node color corresponds to the logarithm of the difference in expression between disease and control samples, where brown means higher expression in disease, grey means no difference, and green stands for higher expression in control. The difference in expression is not shown (white nodes) for targets with very high p-values based on the differential expression analysis (p-value > 0.2).

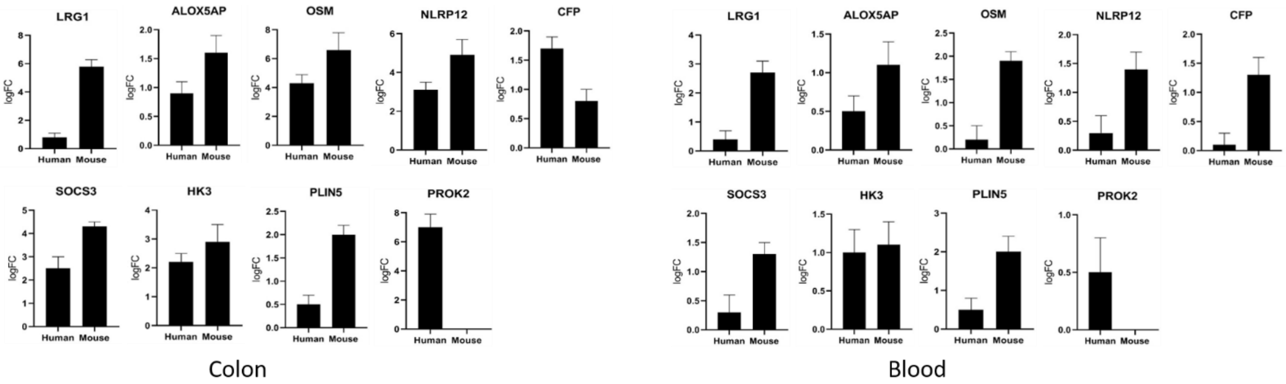


**Supplementary Figure 5.** Validation of differentially expressed genes by real-time qPCR. Using real-time qPCR, the expression of the 9 SDE genes that were identified in our study to be common in the colon and blood of human and mouse UC was evaluated. The real-time qPCR results confirmed the expression patterns. However, PROK2 expression was not detected or lowly detected in the colon and blood of the both mouse and the human control group while in the disease/patients the average cycle quantification value (Cq) was 36, which can explain the high fold change detected for this gene in our RNA-Seq analysis. This gene was labeled as outlier in mouse colon. Overall, lower gene expression was detected in the blood compared to the colon which is the primary site of the disease. It is noteworthy that the UC in the mouse is acute which may explain the higher gene expression in the mouse, while the humans had chronic UC.

**Supplementary Table 1.** Top 10 SDE genes in UC mice colon and blood with average gene expression signal intensity of ≥ 100, padj ≤ 0.05 and FC > 2 for protein-coding genes and lncRNA, and FC > 1.5 for miRNAs.

| Protein coding | | | | | | lncRNA | | | | | | miRNA | | | | | |
|---|---|---|---|---|---|---|---|---|---|---|---|---|---|---|---|---|---|
| Upregulated | | | Downregulated | | | Upregulated | | | Downregulated | | | Upregulated | | | Downregulated | | |
| **Colon** | | | | | | | | | | | | | | | | | |
| **Symbol** | **logFC** | **padj** | **Symbol** | **logFC** | **padj** | **Symbol** | **logFC** | **padj** | **Symbol** | **logFC** | **padj** | **Symbol** | **logFC** | **padj** | **Symbol** | **logFC** | **padj** |
| Saa3 | 16 | 3,10E-15 | Klk15 | -11.3 | 2.0E-17 | 9130221F21Rik | 10.6 | 4.5E-02 | Gm17322 | -6.3 | 1.0E-24 | mir-223-3p | 1.9 | 7.4E-05 | mir-182-5p | -1.4 | 2.9E-05 |
| Reg3b | 15,4 | 1,14E-11 | Mptx2 | -10.3 | 8.4E-14 | E230034D01Rik | 8.7 | 2.3E-04 | 4930552P12Rik | -3.4 | 2.6E-09 | mir-223-5p | 1.7 | 1.8E-11 | mir-192-5p | -1.4 | 1.4E-04 |
| Abca12 | 14,1 | 8,35E-07 | Cyp2c40 | -7.3 | 1.5E-04 | Gm44275 | 7.1 | 7.6E-04 | Gm48832 | -3.3 | 7.8E-15 | mir-224-5p | 1.6 | 7.7E-04 | mir-148a-3p | -1.3 | 3.7E-04 |
| Sptssb | 13,1 | 1,01E-16 | Cyp2c67 | -7.2 | 4.2E-20 | BB123696 | 6.0 | 2.1E-02 | Gm11535 | -3.0 | 1.4E-04 | mir-10b-5p | 1.6 | 8.8E-03 | mir-141-3p | -1.2 | 3.0E-04 |
| Gm49368 | 13 | 0,0007 | Mptx1 | -6.4 | 8.7E-68 | 2310069B03Rik | 4.3 | 3.8E-18 | Gm15401 | -2.9 | 1.9E-16 | mir-10b-3p | 1.6 | 5.9E-03 | mir-210-3p | -1.2 | 7.8E-05 |
| Prss22 | 12,3 | 9,17E-22 | Wnt8b | -6.1 | 1.1E-03 | Gm43189 | 3.6 | 2.0E-02 | Hottip | -2.9 | 3.3E-02 | mir-142a-3p | 1.5 | 4.2E-03 | mir-200a-3p | -1.2 | 5.9E-03 |
| Il6 | 11,8 | 9,96E-09 | Fut9 | -6.0 | 6.1E-27 | Gm19409 | 2.3 | 5.8E-05 | Gm17029 | -2.4 | 4.4E-05 | mir-132-3p | 1.5 | 1.3E-06 | mir-148a-5p | -1.2 | 8.1E-04 |
| Reg3g | 11,5 | 1,35E-07 | Pdzd7 | -5.8 | 5.0E-67 | Gm15987 | 2.2 | 3.3E-03 | Gm10561 | -2.4 | 5.1E-08 | mir-10a-5p | 1.4 | 1.4E-02 | mir-183-5p | -1.2 | 2.5E-04 |
| Pitx2 | 11,5 | 0,0003 | AF529169 | -5.7 | 6.0E-05 | Gm36161 | 2.2 | 9.7E-07 | Has2os | -2.4 | 3.9E-07 | mir-494-3p | 1.4 | 2.9E-05 | mir-200b-3p | -1.2 | 3.8E-03 |
| Ceacam12 | 11,5 | 7,66E-18 | Mettl7a3 | -5.6 | 2.9E-54 | 9130204K15Rik | 2.1 | 5.5E-05 | Hoxa11os | -2.3 | 1.5E-03 | mir-212-5p | 1.3 | 2.0E-09 | mir-194-5p | -1.1 | 9.6E-04 |
| **Blood** | | | | | | | | | | | | | | | | | |
| **Symbol** | **logFC** | **padj** | **Symbol** | **logFC** | **padj** | **Symbol** | **logFC** | **padj** | **Symbol** | **logFC** | **padj** | **Symbol** | **logFC** | **padj** | **Symbol** | **logFC** | **padj** |
| Lipg | 12,1 | 2,7E-04 | Ggnbp1 | -2,6 | 5,5E-05 | Gm7457 | 8,0 | 2,5E-07 | Gm33843 | -6,4 | 0,01 | mir-149-5p | 1.4 | 1.6E-05 | mir-122-5p | -3.5 | 1.6E-05 |
| Gm49368 | 11,7 | 5,5E-22 | P2ry14 | -1,9 | 4,7E-04 | Gm37800 | 7,3 | 1,3E-06 | Gm11934 | -6,2 | 0,02 | mir-223-3p | 1.4 | 3.4E-03 | let-7f-5p | -1.9 | 3.5E-02 |
| Prok2 | 10,6 | 2,6E-05 | Ccr3 | -1,8 | 4,3E-04 | Gm32486 | 7,1 | 4,9E-06 | Gm4673 | -2,9 | 0,0005 | mir-139-3p | 1.0 | 4.4E-02 | mir-126a-3p | -1.4 | 8.9E-04 |
| Saa3 | 10,2 | 4,1E-05 | Gpr174 | -1,7 | 5,3E-05 | Gm33326 | 6,9 | 9,4E-06 | 2900093K20Rik | -1,7 | 0,03 | | | | mir-374b-5p | -1.4 | 1.9E-02 |
| Nos2 | 10,2 | 4,7E-03 | Slc6a19 | -1,3 | 9,5E-04 | Gm45449 | 6,9 | 5,2E-05 | | | | | | | mir-16-5p | -1.1 | 3.5E-02 |
| Itgb2l | 6,2 | 1,3E-03 | Styx | -1,2 | 1,3E-05 | Gm36551 | 6,8 | 1,5E-05 | | | | | | | let-7i-5p | -1.0 | 1.0E-02 |
| Upp1 | 5,2 | 2,1E-20 | Gm42791 | -1,2 | 1,5E-03 | 4933412O06Rik | 6,8 | 1,3E-04 | | | | | | | mir-215-5p | -0.8 | 4.9E-02 |
| Wfdc17 | 4,6 | 2,0E-33 | | | | Gm5441 | 6,6 | 6,4E-05 | | | | | | | | | |
| Lcn2 | 4,5 | 2,8E-04 | | | | E230013L22Rik | 6,5 | 1,6E-04 | | | | | | | | | |
| Steap4 | 4,5 | 6,6E-35 | | | | Gm26912 | 6,0 | 1,4E-03 | | | | | | | | | |
